# Polycystic Ovary Syndrome as an Endogenous Alcoholic Polycystic Ovary Syndrome

**DOI:** 10.1101/143495

**Authors:** IC. Medeiros, JG. Lima

**Affiliations:** Universidade Federal do Rio Grande do Norte, Departamento de Medicina Clínica, Divisão de Endocrinologia – Av. Nilo Peçanha, 620 - Natal/RN, Brazil; Divisão de Gastroenterologia e, Divisão de Endocrinologia – Av. Nilo Peçanha, 620 - Natal/RN, Brazil

**Author notes:** Correspondence to: Ivanildo Coutinho de Medeiros, Rua Hist. Tobias Monteiro, 1863 – Lagoa Nova, Natal-RN, Brazil - CEP 59056-120, Author, Co-author.

## Abstract

Polycystic ovary syndrome (PCOS) is a growing worldwide public health problem that affects millions of women in their reproductive age. Despite being a very common disorder among women, there are still gaps regarding knowledge of disease mechanisms. In this respect, it was recently reported that acetaldehyde (ACD) is endogenously formed during normal ovarian steroidogenesis. The researchers demonstrated that in physiological concentrations ACD caused no detrimental effect on ovarian tissue. Contrariwise, in supraphysiological levels, ACD impairs granulosa cell differentiation, reduces ovulation, and decreases oocyte quality. Gut microbiota of patients with nonalcoholic fatty liver disease (NAFLD) produces significant quantities of endogenous ethanol (EE) and ACD. Because PCOS is closely linked to NAFLD, an ethanol-producing disorder, we hypothesize that it can be an endogenous alcoholic polycystic ovary syndrome (EAPCOS). The main findings of this study were that (i) the odds ratio of having polycystic ovaries is 30-fold greater in alcohol-exposed women than among unexposed controls; (ii) NAFLD/PCOS patients produce gonadotoxic quantities of EE; (iii) NAFLD/PCOS and alcoholic hepatitis individuals share similar liver expression levels of genes regulating high-km ethanol-metabolizing enzymes; (iv) NAFLD/PCOS and alcohol-tolerant drinkers share similar high-capacity to metabolize ethanol in the gut-liver axis; and (v) low blood alcohol concentration (BAC) in NAFLD/PCOS and alcohol-tolerant individuals stem from extensive alcohol degradation in gut-liver axis and significant fecal loss of ethanol. In summary, we provide mechanistic insights supporting the hypothesis that PCOS can be indeed an EAPCOS.

## INTRODUCTION

Polycystic ovary syndrome (PCOS) is the most common cause of anovulatory infertility in women over its reproductive lifetime [1]. Available data indicate that PCOS is closely linked to obesity, insulin resistance, and nonalcoholic fatty liver disease (NAFLD) [2,3]. This suggests that these conditions share a common etiological background.

A striking feature of patients with NAFLD and, by extension with PCOS, is to produce significantly more endogenous ethanol (EE) than controls [4–6]. Accordingly, it is believed that EE plays a critical role in NAFLD development and progression [7–9]. From these observations, we hypothesize that EE also may play a causative role in PCOS pathogenesis.

It is no novelty the intriguing idea about the existence of an endogenous alcoholic disease. Early researchers attempting to validate this hypothesis have provided disappointing results. This discouraged further investigation regarding this matter for several decades. Firstly, because blood-alcohol concentration (BAC) following jejunoileal bypass was significantly lower than in alcoholics, it was concluded that the quantity of EE was insufficient to elicit liver injury [10]. Secondly, since liver histology was normal in a rat model of small intestinal bacterial overgrowth (SIBO), it was inferred that gut bacteria are unable to produce hepatotoxic amounts of ethanol [11]. However, Sprague-Dawley strains only develop steatohepatitis and fibrosis 12 to14 weeks after study initiation. And in this study, the rodents were sacrificed very early, around the 4th to 5th weeks of experimentation [12]. Therefore, as this mouse model recapitulates the histopathological spectrum of human alcoholic liver disease, we postulate that it can also cause polycystic ovaries.

Thus, the main focus of this study was to provide a mechanistic explanation of how PCOS may be an endogenous alcoholic polycystic ovary syndrome (EAPCOS).

### Ethanol is a prodrug

As a prodrug, ethanol requires conversion to ACD in order to exert its cytotoxic properties [13,14]. This pharmacokinetic characteristic of ethanol is of paramount importance for a mechanistic understanding of the EAPCOS hypothesis. The clear and obvious implication of this is that in a scenario of extensive presystemic catabolism of ethanol, BACs can be negligible or even undetectable. The finding that intravenous infusion of cirrhotogenic amounts of ethanol (57.6 to 115.2 g/d) in alcohol-tolerant drinkers induces only modest BACs (1.5 to 4.5 mg/dL) supports this notion [15].

### Gastrointestinal production and degradation of ethanol

Gastrointestinal microbiota of healthy subjects produces nontoxic amounts of EE from luminal dietary carbohydrates. Next, EE is absorbed and metabolized in the liver to ACD, which in turn is oxidized to non-toxic amounts of acetate [16]. In contrast, in SIBO-related conditions such as NAFLD/PCOS [17][18] and malabsorption syndromes [19], gut production of EE is significantly greater than in controls [4–6]. Agreeing with this, gut/fecal concentrations of EE in these cases are proportionally equal to or even greater than those obtained after moderate drinking [19–21]As ethanol is formed within a dysbiotic gut, it is converted to ACD in a dose- and concentration-dependent manner [11]. In this setting, ACD-producing alcohol dehydrogenase (ADH) activity is higher than that of ACD-oxidizing aldehyde dehydrogenase (ALDH) [22]. The net result is ACD build-up coupled with low BACs. However, in the auto-brewery syndrome, massive production of EE exceeds gut-liver axis ability to clear alcohol from circulation. As a consequence, BAC may reach 250-350 mg/dL [23].

Blind-loop contents of rat jejunum converts ethanol to ACD at a rate of 1.99 μM/min ● mL under aerobic conditions [11]. If applicable to patients with SIBO and small bowel contents around 2000 mL, it is estimated that everyone can convert ~528 g of EE into ACD daily. Experimentally it has been demonstrated that extrahepatic ACD is 30 to 330-fold more hepatotoxic than that formed intrahepatically [24,25]. Extrapolating this to humans, 0.18-2 g of EE will provide an amount of ACD as hepatotoxic as that generated intrahepatically from a cirrhotogenic dose of 60 g ethanol [26].

BACs found in NAFLD/PCOS as well as in alcohol-tolerant subjects are consistently low [15,27]. In this scenario, for maintaining a steady-state BAC of 7.14 mg/dL [27] NAFLD/PCOS patients would need a 24-h continuous intravenous infusion of ethanol at a rate ~9.5 g/h (228 g/day) [28].

Lastly, a seminal study showed up-regulation of all genes involved in ethanol metabolism in nonalcoholic steatohepatitis livers [5]. Importantly, these genes encode enzymes those maximal catalytic activities are at high ethanol concentrations [29]. Even more noteworthy is the finding that hepatic expression of ethanol-metabolizing genes in NAFLD/PCOS is similar to that of alcoholic hepatitis [7].

### Ethanol pharmacokinetics in NAFLD/PCOS is similar to that of alcohol-tolerant individuals

Blood-alcohol elimination rate may be 3-fold to 4-fold higher in alcohol-tolerant than in healthy individuals and social drinkers [29]. Causative factors include induction of high-Km alcohol-metabolizing enzymes by ethanol, insulin resistance, ketone bodies, unsaturated fatty acids [30], iron overload [31], hyperglycemia [32], and gut microbial degradation of ethanol [11]. Additionally, genetic polymorphisms of alcohol-metabolizing enzymes play an essential role in the development of this so-called alcohol tolerance/adaptation process [33,34]. In this setting, instead of metabolizing ethanol in milligram levels through ADH1 activity, the individual starts doing this at tens of grams scale via CYP2E1, catalase, ADH3, and ADH4 [30,35–37]For example, for maintaining a steady-state BAC between 1.5 to 4.5 mg/dL, alcohol-tolerant individuals require a continuous intravenous infusion of ethanol at a rate equivalent to 57.6 to 115.2 g/d [15].

Interestingly, and probably not by chance, all these risk factors for increased blood-alcohol clearance are also present in NAFLD/PCOS [5,38–40]. For this reason, it is intuitive to infer that patients with NAFLD/PCOS metabolize exogenous/endogenous ethanol similarly to alcohol-tolerant individuals (Figure). In line with this argument, (i) aberrant microbiota found in NAFLD as well as in alcoholics consists of both ethanol-producing and -degrading bacteria [4,41,42], (ii) gut production and degradation of ethanol are dose-dependent and simultaneous processes that prevent high BAC [11,19,43,44], and (iii) gut concentrations of ACD reach mutagenic values (49-87 μM) in rat blind loops [11]. Furthermore, impaired gastrointestinal absorption [45–48] and binding/entrapment of alcohol in food constituents certainly contribute to low BACs [47,49]. Notably, fecal loss of ethanol reaches concentration values up to 50-fold greater than BACs found in NAFLD [19][27].

**Figure 1.**
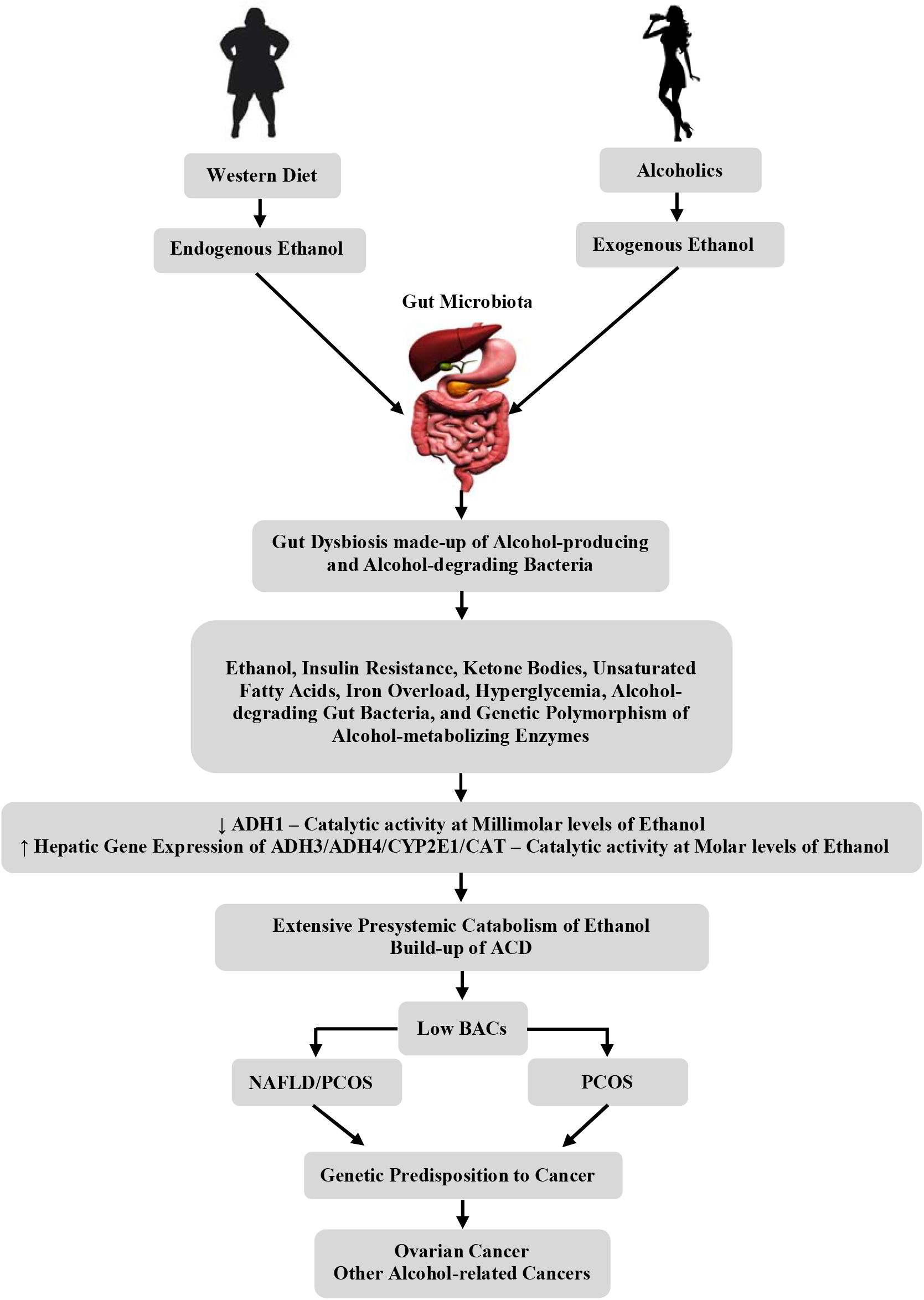
Evidence that NAFLD/PCOS patients metabolize etanol similarly to Alcohol-tolerant individuals. Chronic exposure either to endogenous or exogenous high ethanol concentrations leads to a gut dysbiosis made-up of alcohol-producing and - degrading organisms. Then, a number of metabolic alterations switch alcohol metabolism from milligram scale to hundreds of grams per day. This extensive presystemic clearance of ethanol causes ACD build-up, which in turn leads to the consistently low BACs. Lastly, in genetically predisposed women carcinogenic ACD can lead to ovarian cancer and other alcohol-related neoplasms. Km = Michaelis Constant; ADH = Alcohol dehydrogenase; CYP2E1 = Cytochrome P-450 2-E1; CAT = Catalase; EAPCOS = Endogenous Alcoholic Polycystic Ovary Syndrome; PCOS = Polycystic Ovary Syndrome; ACD = Acetaldehyde;

### Calculating EE production from a standard human pharmacokinetic model

One study showed that average BAC of twenty patients with NAFLD was 7.14 mg/dL after a 12h overnight fast [27]. Assuming that ethanol elimination rate of these individuals was 20 mg/dL/h and that its mean height was 1.74 m, we estimate the amount of EE produced by each patient. To achieve this, we use a validated physiologically-based model of alcohol metabolism, which has been described in detail elsewhere [28]. According to our calculations, each patient produced 161.49 g of EE after a 12-h overnight fast. By extrapolating the data to three equicaloric meals, the daily production of EE should reach 484 g. Because EE undergoes extensive conversion to ACD in the gut-liver axis, estimated total circulating alcohol burden is only 4.6 g. This leads to the characteristically low BAC found in these patients (see [28] for equations and calculation procedures).

### Ovariotoxicity of ethanol

It has been shown that endogenous ACD is formed as a byproduct during normal ovarian steroidogenesis [50]. In this situation, there is no ovotoxicity because it converted to acetate by ovarian ALDH. On the contrary, in supraphysiological concentrations, ACD disrupts the differentiation of granulosa cells, reduces ovulation and lowers oocyte quality [50]. Acute as well as chronic exposure to ethanol inhibits ovarian steroidogenic acute regulatory protein (StAR), which plays a crucial role in gonadotropin-stimulated gonadal hormone production [51,52]. This leads to an increase in the number of the corpora lutea, atretic follicles, regression of theca antral follicles, vacuolation/fat deposition in granulosa, theca, and interstitial cells [53]. As a result, alcohol exposure entails disruption of puberty and menstrual cycling as well as a hormonal imbalance in pre and postmenopausal women [54].

Agreeing with these observations, the odds ratio of having PCOS is about 30-fold higher among Chinese women consuming alcohol than in matched controls [55]. This substantial discrepancy is because 30% to 40% of Asian individuals carry out a defective ALDH2 enzyme, providing the accumulation of very high quantities of ACD [56,57]. Moreover, subgroup analyses of multiple subpopulations suggest a positive relationship between heavy alcohol intake and ovarian cancer [58]. Supporting argument for this epidemiological connection comes from the finding of elevated concentrations of ACD in ovarian cancer tissues [59].

### Other endogenous and exogenous risk factors for PCOS

According to the data presented here, ethanol itself recapitulates the entire phenotype of PCOS. However, other endogenous/exogenous toxins may be implicated in its pathogenesis. These include dicarbonyl compounds [60–62], advanced glycation end products (AGEs) [63,64], advanced lipid peroxidation end products (ALEs) [65,66] and nitric oxide radicals [67]. Furthermore, it is possible that epigenetic changes and aberrant microRNA (miRNA) may play a relevant role in PCOS development [68,69]. Considering that NAFLD/PCOS is an alcohol-producing disorder, we postulate that even alcohol-abstinent women are at increased risk of developing ovarian cancer. This event would be particularly likely among NAFLD/PCOS patients carrying genetic susceptibility to both sporadic and hereditary ovarian neoplasms [70,71]. Likewise, NAFLD/PCOS women carrying functional polymorphisms in ethanol-metabolizing genes may be at highest risk for developing ovarian cancer and other alcohol-related neoplasms [72,73].

## Conclusion

In short, our data reconcile the apparently contradictory association between exposure to large amounts of ethanol concurrently with negligible BACs [15]. The EAPCOS hypothesis contains several limitations, including the fact that most of the evidence comes from uncontrolled observational studies. Nevertheless, it provides sufficient evidence justifying the existence of an EAPCOS. Lastly, if confirmed by further studies, our hypothesis may contribute to novel therapeutic and preventive strategies for disease management and control.

## Author’s roles

I.C.M. elaborated the study’s hypothesis and purpose, performed the literature search, wrote and critically reviewed and approved the final version of the paper; J.G.L. contributed to the study design, figure elaboration, paper editing, and critically reviewed and approved the final version of the manuscript.

